# Structural insights into physiological activation and antagonism of melanin-concentrating hormone receptor MCHR1

**DOI:** 10.1101/2024.03.12.584619

**Authors:** Xiaofan Ye, Guibing Liu, Xiu Li, Binbin He, Yuyong Tao, Haiping Liu, Weimin Gong

## Abstract

Melanin-concentrating hormone (MCH) is a 19-amino-acid neuropeptide playing crucial roles in energy homeostasis, sleep, and various physiological processes. It acts through two G protein-coupled receptors, MCHR1 and MCHR2, with MCHR1 being universally present in mammals and a potential target for treating metabolic and mental health conditions. However, drug development efforts have been impeded by the lack of structural information. Here, we present the cryo-EM structures of MCHR1 in its active state complexed with MCH and G_i1_, as well as in its inactive state bound to a selective antagonist SNAP-94847. Structural and mutagenesis analyses disclosed the recognition mechanisms for both MCH and SNAP-94847, the activation mechanism and antagonism of MCHR1, and the determinants of ligand specificity. These findings are expected to accelerate the development of better drugs targeting the MCH system.

## Introduction

Melanin-concentrating hormone (MCH) was initially identified in salmon as a cyclic heptadecapeptide involved in the regulation of pigmentation^1^. However, mammalian MCH does not govern skin color but instead plays a crucial role in regulation of feeding behavior and energy homeostasis^2–4^. In rodents and humans, MCH is a neuropeptide expressed in hypothalamus and zona incerta^5^. Experiments using mouse models found that MCH expression is upregulated in fasting conditions, and administration of MCH stimulates food intake and promotes body weight gain^6,7^. MCH-knockout mice manifest reduced appetite, become emaciated, and exhibit increased energy expenditure and locomotor activity^8^. Additionally, MCH significantly influences sleep by facilitating the occurrence of slow wave sleep (SWS) and rapid eye movement sleep (REMS) through inhibiting the wakefulness-inducing neurotransmitter system^9^, thus establishing its classification as a neurotransmitter involved in sleep regulation^10^. Furthermore, MCH exerts its influence on various behavioral and physiological functions such as reward processing, learning, olfaction, anxiety, and nociception^11–13^.

Currently, two high-affinity receptors for MCH, MCHR1 and MCHR2, have been identified in human^4^. Both receptors belong to the G protein-coupled receptor (GPCR) superfamily, with MCHR1 being common to all mammals and well-characterized in terms of its role in physiological processes. MCHR1 is primarily expressed in the central nervous system^14^, but can also be expressed in brown adipose tissue^15^, and is coupled by G_i_, G_o_, and G_q_^16^. The primary function of the MCH-MCHR1 system is in regulation of feeding behavior and energy homeostasis^4^. Studies have shown that knockout of MCHR1 in mice leads to a lean phenotype^17^, and naturally occurring MCHR1 mutations in humans have been found to disrupt MCHR1 signaling, resulting in underweight phenotypes^18^. Additionally, the MCH-MCHR1 system is involved in various neuronal functions, such as sleep, mood, and learning^19–21^. Administration of selective MCHR1 antagonists has been shown to induce antianxiety and antidepressant effects in rodents^22–24^, and MCHR1 knockout mice also display an antianxiety phenotype^25^. The MCH-MCHR1 system has also been implicated in regulating primary cilia growth and function^26–28^. MCHR1 is considered a potential therapeutic target for various diseases, including obesity, sleep disorders, anxiety disorders, schizophrenia, and Alzheimer’s disease^29–32^. Numerous MCHR1 antagonists, such as SNAP-7941, AMG-076, NGD-4715, AZD1979, SNAP-94847 and RGH-076, have been developed and some have shown therapeutic effects in clinical studies^29,33^. Given the complex role of MCHR1 in both physiological and pathological conditions, further research on the modes of interaction between MCHR1 and its activating or inhibiting ligands is of great importance.

In this study, we present cryo-EM structures of the active-state MCHR1 in complex with MCH and G_i1_ and the inactive-state MCHR1 bound to a high-affinity and highly selective antagonist SNAP-94847^34^. Through a combination of structural comparison and functional assays, we elucidate the recognition mechanisms of MCH and SNAP-94847 by MCHR1 and unveil the activation mechanism and antagonism of MCHR1. These findings are expected to facilitate the discovery of better drugs targeting the MCH system.

## Results

### Structure of the MCH-MCHR1-G_i1_ complex

To acquire the MCH-MCHR1-G_i1_ complex, we co-expressed human MCHR1 and the three subunits of G_i1_ in Sf9 insect cells. A NanoBiT tethering strategy was applied to promote complex formation by increasing local concentration of G proteins^35^. Additionally, a single-chain Fab fragment (scFv16) was utilized to stabilize the complex^36^. The complex was purified in the presence of synthetic MCH peptide and subjected to cryo-EM analysis. The structure was determined in multiple states, of which the best reach the resolution of 2.61 Å (Fig. 1a, Extended Data Figs. 1 and 2, and Supplementary Table 1). These states are primarily distinguished by the relative orientation between MCHR1 and the G_i1_ heterotrimer. Two states (T1 and T2) exhibit tighter receptor-G protein contact over the other two states (L1 and L2). The tight conformers accounts for 53.4% of the particles that passed the screening cycles, while the loose conformers accounts for the remaining 46.6% particles (Extended Data Fig. 1b). G_i_-coupling modes and the conformations of the Gα subunit are different in these structures. Specifically, the first intracellular loop (ICL1) of MCHR1 adopts distinct conformations in different states (Extended Data Fig. 2e). The tight conformers share a similar ICL1, which makes a close contact with G_i1_ at the αN-Gβ interface (Extended Data Fig. 2f). In contrast, in the loose conformers, ICL1 only makes a loose contact with Gβ (Extended Data Fig. 2g). Moreover, when the structures are aligned by the receptor, with the carboxyl terminus of α5 helix almost fixed, we observed a sequential displacement of the αN helix, which undergoes a clockwise rotation from the L2 state to the T2 state (Extended Data Fig. 2h). Despite the differences, MCHR1 in all these conformations is in a typical fully active state. These subtle differences may reflect limited flexibility of the nucleotide-free GPCR-G protein complexes.

**Fig. 1.**
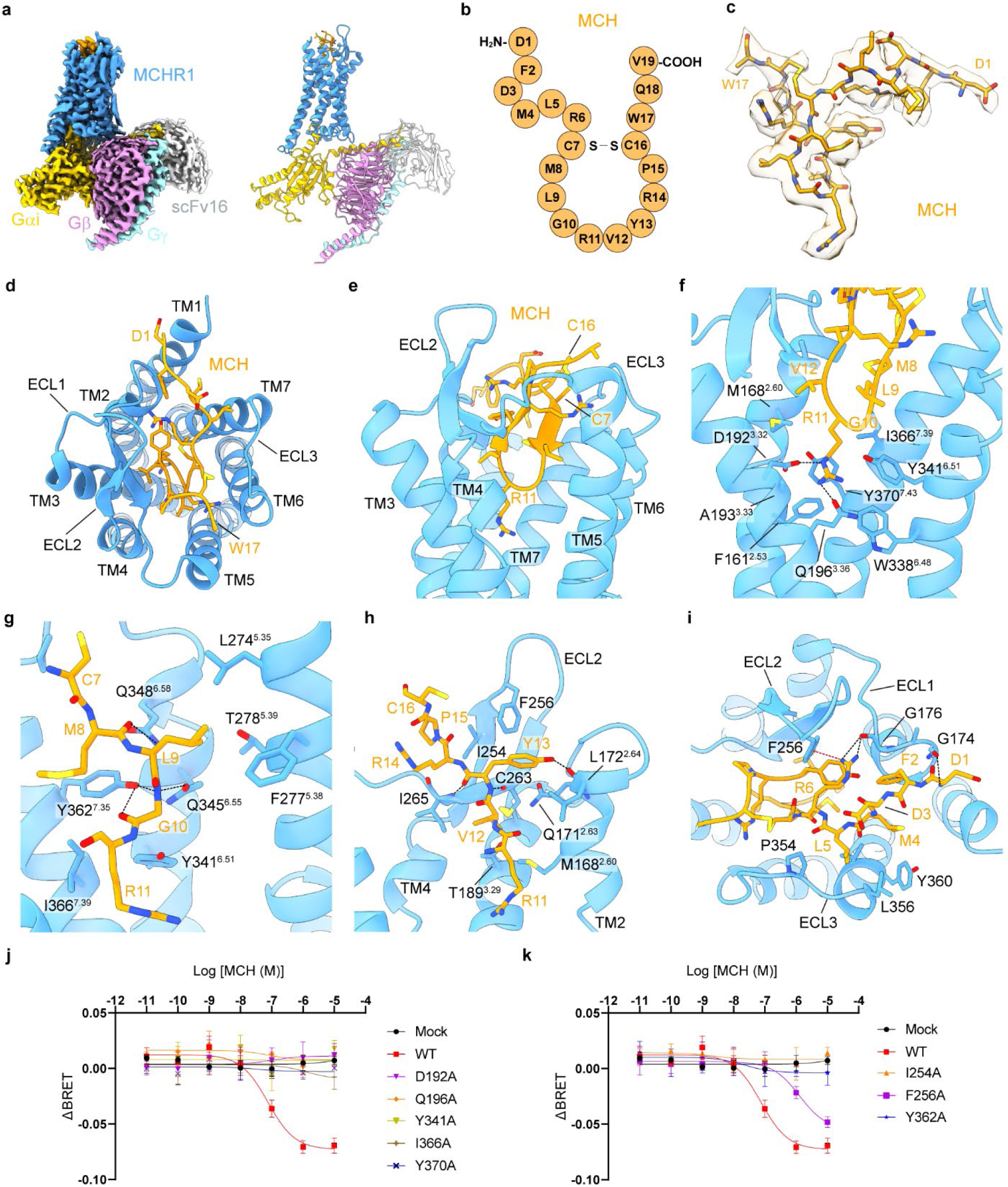
Cryo-EM structure of the MCH-MCHR1-G_i1_ complex. **a**, EM map and model of MCH-MCHR1-G_i1_ complex in T1 state. b, Sequence of human MCH. c, Density map of MCH in T1 state. d, Extracellular view of MCH-MCHR1-G_i1_ complex in T1 state. e, The β-hairpin of MCH and the binding pocket. f, Interactions between R11 and MCHR1. The hydrogen bonds are depicted as black dashed lines. g, Interactions between the N-terminal fragment of the β-hairpin and MCHR1. h, Interactions between the C-terminal fragment of the β-hairpin and MCHR1. i, Interactions between the N-terminus of MCH and MCHR1. The cation-π interaction between R6 and F256 is depicted as a red dashed line. j-k, G_i_-dissociation curves of MCHR1 mutants. Data are shown as means ± SEM from three independent experiments.

### Recognition of MCH by MCHR1

Human MCH is a 19-amino-acid peptide cyclized by the disulfide bond between **C7** and **C16** (Fig. 1b). Despite the conformational variations of the MCH-MCHR1-G_i1_ complexes, the binding pose of MCH in MCHR1 among different states remains the same (Extended Data Fig. 3a). The density from **F2** to **W17** is well-defined in the EM maps except for the side chain of **W17** (Fig. 1c and Extended Data Fig. 2a-d). MCH binds in a large pocket formed by the extracellular side portions of the transmembrane helices (TMs) 2, 3, 5, 6, and 7, as well as extracellular loops (ECLs) 1, 2, and 3 (Fig. 1d). Notably, ECL2 also forms a pair of antiparallel β-strands as observed in other peptide receptors.

**C7**-**C16** of MCH is cyclized and forms a short β-hairpin, which inserts into the central pocket formed by transmembrane helices (Fig. 1e). The most notable feature is that **R11**, which locates at the distal end of the β-hairpin, extends deeply into the pocket. The side chain of **R11** is stabilized by hydrogen bonds with D192^3.32^ and Q196^3.36^ (superscripts represent Ballesteros-Weinstein nomenclature), as well as hydrophobic interactions with F161^2.53^, M168^2.60^, A193^3.33^, W338^6.48^, Y341^6.51^, I366^7.39^, and Y370^7.43^ (Fig. 1f).

Mutations of D192^3.32^, Q196^3.36^, Y341^6.51^, I366^7.39^, and Y370^7.43^ into alanine dramatically impaired MCH-induced G_i_ dissociation in MCHR1-expressing HEK293T cells (Fig. 1j and Extended Data Fig. 3b), indicating the crucial roles of these residues in MCHR1 activation. In addition, M168^2.60^A also mildly impaired MCHR1 activation (Extended Data Fig. 3c). On the N-terminal side of the β-hairpin, the backbone atoms of MCH forms hydrogen bonds with the sidechains of Q345^6.55^, Q348^6.58^, and Y362^7.35^ (Fig. 1g). Mutation of Y362^7.35^ into alanine dramatically impaired MCH-induced G_i_ dissociation (Fig. 2k), while mutation of Q345^6.55^ exhibited a mild effect (Extended Data Fig. 3d). On the C-terminal side of the β-hairpin, **Y13** interacts with the second β-strand of ECL2 through backbone hydrogen bonds, while its sidechain also forms a hydrogen bond with the backbone carbonyl of Q171^2.63^ (Fig. 1h). In addition, residues on the first strand of ECL2 also establish contacts with MCH, with F256 acting like a wedge that prevents the release of MCH (Fig. 1h). MCH exhibited decreased potency to F256A mutant in G_i_-dissociation assay (Fig. 1k). Unexpectedly, I254A mutation almost abolished MCH-induced G_i_ dissociation (Fig. 1k), possibly due to improper folding of ECL2.

**Fig. 2.**
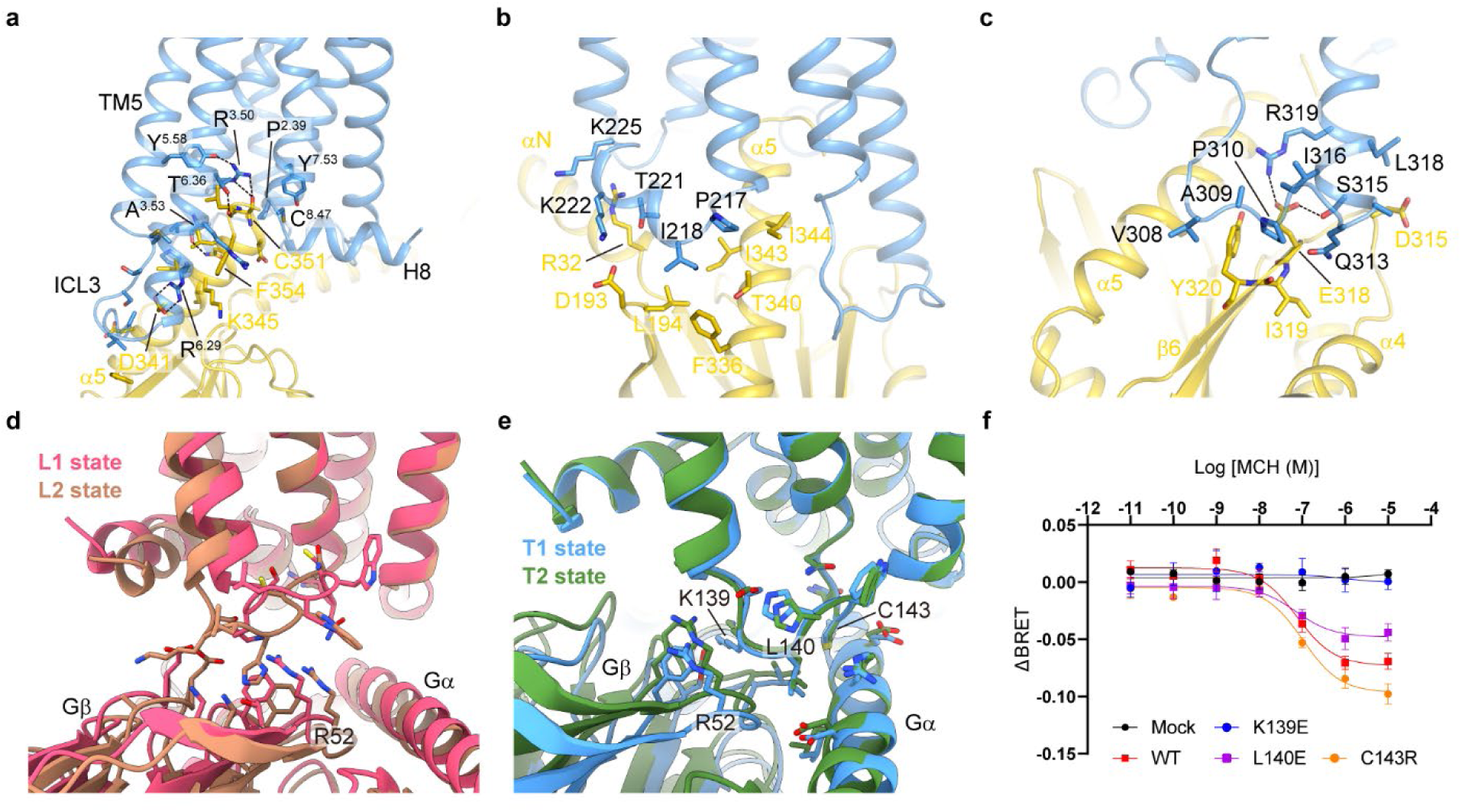
G_i_ coupling of MCHR1. **a**, Interactions between MCHR1 and the α5 helix of Gα_i1_. The hydrogen bonds are depicted as black dashed lines. **b**, Interactions between ICL2 of MCHR1 and the αN-α5 hydrophobic patch. **c**, Interactions between ICL3 of MCHR1 and Gα_i1_. **d**, The contact between ICL1 and G_i1_ in the loose states. **e**, The contact between ICL1 and G_i1_ in the tight states. **f**, Effects of ICL1 mutations determined by G_i_-dissociation assay. Data are shown as means ± SEM from three independent experiments.

The N-terminus of MCH binds in a superficial sub-pocket mainly by interactions with extracellular loops of the receptor (Fig. 1i). The amino group of **D1** forms a hydrogen bond with backbone carbonyl of G174^ECL^^1^. **R6** forms a hydrogen bond with backbone carbonyl of G176^ECL^^1^ and a cation-π interaction with F256^ECL^^2^ (Fig. 1i). **F2** and **L5** are tightly packed with ECL1 and ECL3, respectively. It is noteworthy that MCH_6-17_ exhibits reduced affinity with MCHR1 compared to full-length MCH^37^, indicating the role of the N-terminus in increasing the binding affinity.

### G_i_ coupling of MCHR1

Resembling other GPCR-G protein complexes, the primary interface between MCHR1 and G_i1_ is also formed by the intracellular cavity of MCHR1 and the α5 helix of Gα_i1_. The C terminus of α5 helix inserts into a cavity formed by residues from TM3, ICL3 and TM6, as well as P147^2.39^, Y297^5.58^, Y380^7.53^, and C384^8.47^ (Fig. 2a). Notably, R210^3.50^, A213^3.53^, R319^6.29^, and T326^6.36^ form polar interactions with residues of the α5 helix. ICL2 of MCHR1 forms an ordered short helix and makes a contact with the αN-α5 hydrophobic patch (Fig. 2b). Moreover, ICL3 of MCHR1 is mostly ordered in the structures. Together with the intracellular end of TM6, ICL3 forms an additional interface with Gα subunit at the β6 strand and the α4-β6 loop (Fig. 2c), where S315 and R319 forms polar interactions with E318 of Gα.

Interactions between ICL1 and G protein were less observed previously. In the loose conformations of the MCH-MCHR1-G_i1_ complex, ICL1 only makes a loose contact with the surface of Gβ (Fig. 2d). However, in the tight conformers, K139, L140, and C143 of ICL1 make close contacts with G_i1_ at the αN-Gβ interface (Fig. 2e). Specifically, the sidechain of L140 inserts into the crevice between αN helix and Gβ. To investigate the role of ICL1 in MCHR1 activation, we introduced mutations at ICL1 and measured MCH-induced G_i_ dissociation in MCHR1-expressing HEK293T cells. Among the three sites mutated, L140E and C143R had little effects, while K139E dramatically impaired activity of MCHR1 (Fig. 2f and Extended Data Fig. 3b), suggesting a potential role of ICL1 in G_i_ activation.

### Structure of antagonist-bound MCHR1

MCHR1 antagonists are potential drugs for metabolic and psychiatric diseases. To understand the antagonism of MCHR1, we tried to solve the structure of the inactive-state MCHR1 bound to a selective antagonist SNAP-94847. To this end, we applied a previously described strategy^38^ to determine this structure by cryo-EM. We engineered the human MCHR1 by inserting mBRIL between TM5 and TM6 to replace the ICL3 in a rigid fashion and fused a K3 helix together with an ALFA tag to H8 of MCHR1. Besides, we introduced the following components, an anti-BRIL Fab (Fab1B3) for engaging mBRIL and a bivalent glue molecule containing anti-Fab and anti-ALFA nanobodies (NbFab and NbALFA) for conjugation of anti-BRIL Fab and H8-K3-ALFA. The engineered construct was purified in the presence of SNAP-94847 and incubated with Fab1B3 and the glue molecule in vitro to obtain the antagonist-bound MCHR1. Single-particle cryo-EM resulted in the three-dimensional reconstruction of SNAP-94847-MCHR1 at 3.33 Å resolution (Fig. 3a and S1 state in Extended Data Fig. 4b). Another conformation with a lower resolution was also observed (S2 state in Extended Data Fig. 4b). Despite differences in overall conformation of the entire complex, the MCHR1 part remains almost the same in these two states (Extended Data Fig. 5a). Except for the flexible cellular loops, the cryo-EM map allowed unambiguous assignment of the majority of MCHR1 (Fig. 3b and Extended Data Fig. 4g). Additional density was observed inside the transmembrane helices and modeled as SNAP-94847 (Fig. 3c).

**Fig. 3.**
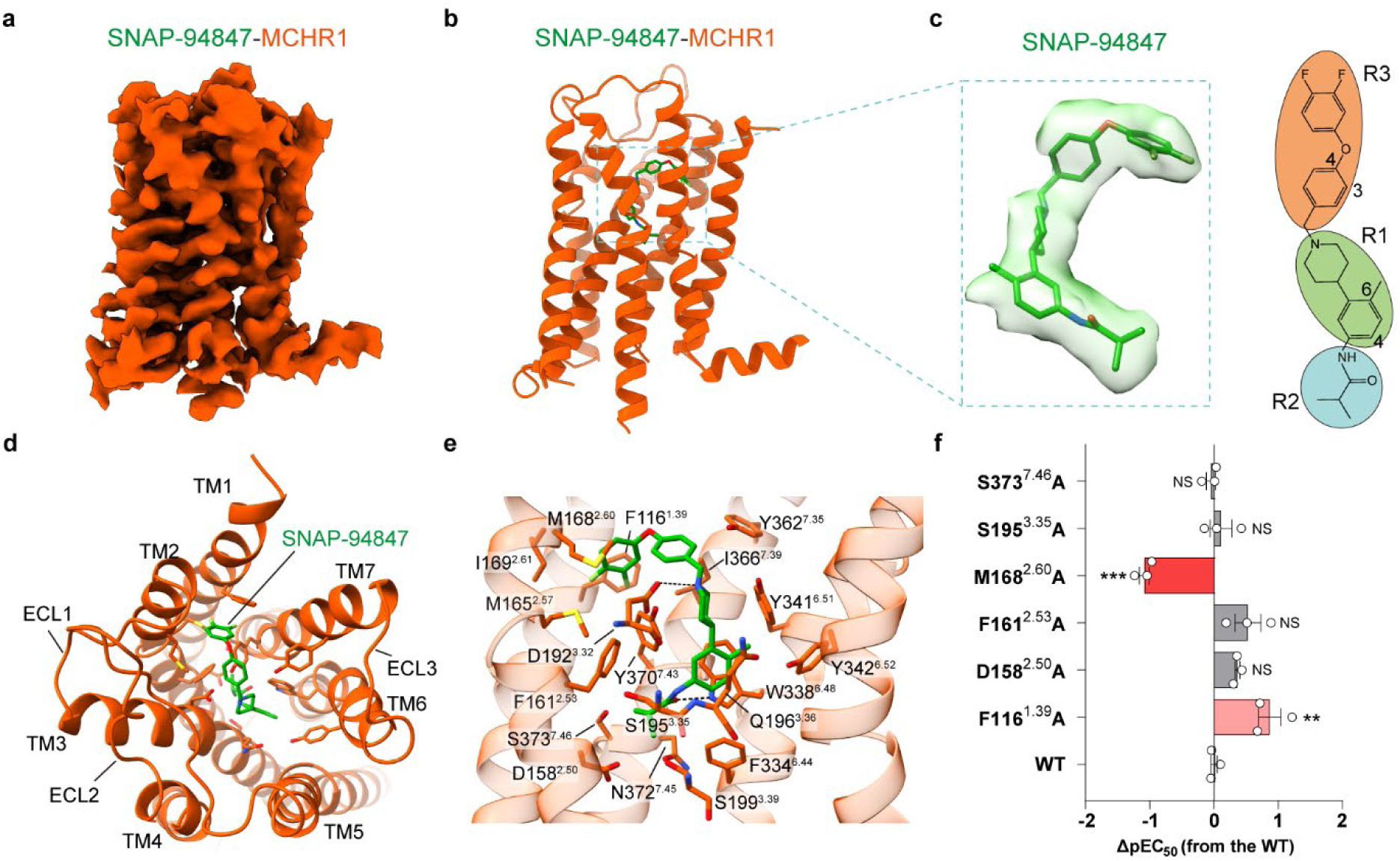
Cryo-EM structure of antagonist-bound MCHR1. **a**, Cryo-EM map of antagonist-bound MCHR1 in the S1 state. **b**, Model of antagonist-bound MCHR1. SNAP-94847 is shown as green sticks. **c**, Density map and molecular structure of SNAP-94847. **d**, Top view of the structure. **e**, Interactions between SNAP-94847 and MCHR1. The ionic interactions and hydrogen bonds are depicted as black dashed lines. **f**, G_i_-dissociation assay results of MCHR1 mutants in response to SNAP-94847. ΔpEC_50_ of each mutant is compared to WT using one-way ANOVA with Dunnett’s multiple comparisons. \*\**P* < 0.01, \*\*\**P* < 0.001. NS, no significant difference.

The overall structure of the inactive MCHR1 resembles that of the active MCHR1 (Extended Data Fig. 5b). We observed the hallmark of GPCR inactive conformation in the SNAP-94847-MCHR1 structure, the inward movement of the intracellular end of TM6 (Extended Data Fig. 5b). This rearrangement triggers the closure of the cytoplasmic pocket to prevent receptor coupling with downstream effectors, leading to receptor inactivation.

### Antagonist binding of MCHR1

The antagonist SNAP-94847 can be divided into three functional groups, a 4-(2-methylphenyl)piperidine scaffold (R1), an isobutyramido group (R2), and a 4-(3,4-difluorophenoxy)benzyl group (R3) (Fig. 3c). It binds into a hydrophobic pocket surrounded by TMs 1, 2, 3, 6, and 7 (Fig. 3d). Unlike the agonist MCH, the antagonist SNAP-94847 binds deeper within the 7TM domain and its R2 group extends inward by about 7.5 Å distance compared with **R11** of MCH (Extended Data Fig. 5c). Specifically, the majority of the R1 group makes tight hydrophobic interactions with Q196^3.36^, F334^6.44^, W338^6.48^, Y341^6.51^, Y342^6.52^, and Y370^7.43^, whereas the quaternary amine forms a strong ion-ion interaction with D192^3.32^ (Fig. 3e). The R2 group is buried in a compact sub-pocket deep inside the 7TM domain, composed of D158^2.50^, F161^2.53^, S195^3.35^, S199^3.39^, F334^6.44^, W338^6.48^, N372^7.45^, and S373^7.46^. The interaction is stabilized by a hydrogen bond between the carbonyl oxygen of the R3 group and W338^6.48^ (Fig. 3e). Besides, the R3 group of SNAP-94847 forms hydrophobic interactions with F116^1.39^, M165^2.57^, M168^2.60^, I169^2.61^, Y362^7.35^, I366^7.39^, and Y370^7.43^ (Fig. 3e).

Interaction sites are partially identical for SNAP-94847 and MCH, such as D192^3.32^, Q196^3.36^, Y341^6.51^, I366^7.39^, Y370^7.43^ (Fig. 1f and Fig. 3e). SNAP-94847 competitively interacts with these sites and block MCH binding. In addition, in mutagenesis studies, mutation of M168^2.60^ to alanine significantly reduced potency of SNAP-94847 (Fig. 3f and Extended Data Fig. 5d, e), suggesting a crucial role of M168^2.60^ in determining the binding affinity with antagonists. Unexpectedly, mutation of F116^1.39^ to alanine significantly increased potency of SNAP-94847. Mutation of the phenylalanine to alanine may contribute to better accommodation of the bulky 3,4-difluorophenoxy group.

### Activation mechanism of MCHR1

This study presents the structures of the endogenous agonist-bound active and antagonist-bound inactive forms of MCHR1, offering insights into the mechanisms of receptor activation. The structures of the MCH-MCHR1-G_i1_ complex and the SNAP-94847-MCHR1 complex were superimposed, revealing significant differences in both the extracellular and intracellular regions. Binding of MCH prompts an inward shift at the extracellular ends of TMs 2, 3, and 4, as well as ECL2, resulting in contraction of the orthosteric ligand-binding pocket (Fig. 4a). On the intracellular side, there is a notable outward movement of TM6, which facilitates the coupling with G protein (Fig.4b).

**Fig. 4.**
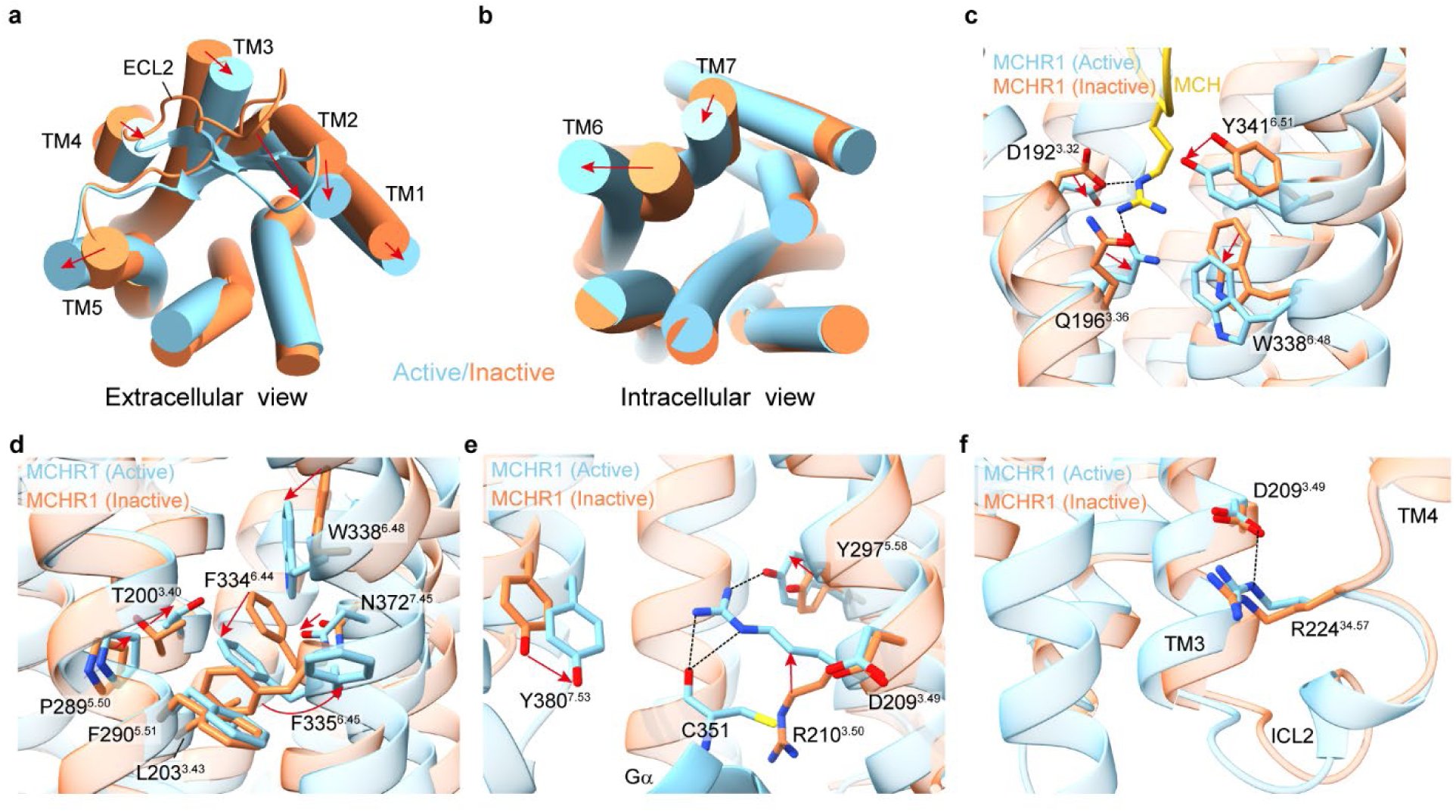
Activation of MCHR1. **a**, Superposition of active MCHR1 (T1 state) and inactive MCHR1 (S1 state) in the extracellular view. Transmembrane helices (TMs) are shown as cylinders. The movement of TMs are indicated by red arrows. **b**, Intracellular view of the superposed structures. **c**, Comparison of the ligand-binding pocket. **d**, Rearrangement of hydrophobic packing. **e**, Comparison of the DRY motif. **f**, Association between D^3.49^ and ICL2. The ionic interactions and hydrogen bonds are depicted as black dashed lines.

Upon activation, extensive interactions between MCH and MCHR1 drives a remarkable downward shift of the indole ring of W338^6.48^ (Fig. 4c). W338^6.48^ induces re-packing of T200^3.40^, L203^3.43^, P289^5.50^, F290^5.51^, F334^6.44^, F335^6.45^, and N372^7.45^, initiating the outward movement of TM6 at the cytoplasmic end (Fig. 4d). Notably, F335^6.45^ undergoes a drastic rotation from inside of the 7TM domain to the outside.

MCHR1 has a conserved D^3.49^R^3.50^Y^3.51^ motif. However, in the inactive structure, R210^3.50^ is not locked by D209^3.49^ as observed in many inactive class A GPCR structures (Fig. 4e). In the active state, R210^3.50^ forms hydrogen bonds with Y297^5.58^ and the backbone carbonyl of C351 from Gα_i1_, facilitating the adoption of the active conformation (Fig. 4e). Furthermore, D209^3.49^ of MCHR1 forms an ionic interaction with R224^34.57^ of ICL2 in both states, potentially contributing to the stabilization of ICL2 (Fig. 4f).

## Discussion

In this study, we identified different conformational states of the MCH-bound MCHR1-G_i1_ complex. Recently, two different conformational states of the acetylcholine-bound M_2_ muscarinic acetylcholine receptor (M_2_R)-G_oA_ complex^39^ were also identified compared with the single state of M_2_R-G_oA_ in complex with the more potent agonist iperoxo^40^. Structural comparison revealed that the orientation of Gα subunit differs in two states (S1 and S2) of the acetylcholine-M_2_R-G_oA_ complex, with a smaller αN-α5 angle in the S2 state than in the S1 state (Extended Data Fig. 6a). NMR studies on the conformational dynamics of M_2_R supported that iperoxo is more efficacious in stabilizing a uniform nucleotide-free M_2_R-G_oA_ signaling complex than acetylcholine. Similarly, in our study, the tight conformers of MCHR1-G_i1_ complex exhibit a smaller αN-α5 angle than the loose conformers (Extended Data Fig. 6b). The balance of different conformations is potentially associated with receptor activation by different agonists.

MCHR2 is an additional functional MCH receptor in humans and may play a role in MCH-related physiological functions. Although it shares 37% sequence identity with MCHR1, it is not yet clear if MCHR2 utilizes the same ligand recognition mechanism. Alignment of 33 residues responsible for MCH recognition by MCHR1 has shed light on this matter. In MCHR2, 22 of these residues are similar to those in MCHR1 (Fig. 5a). In addition, the non-conserved residues are predominantly located in peripheral areas and make fewer contacts with MCH (Fig. 5b). This suggests that the MCH-binding mechanisms of the two receptors are relatively conserved.

**Fig. 5.**
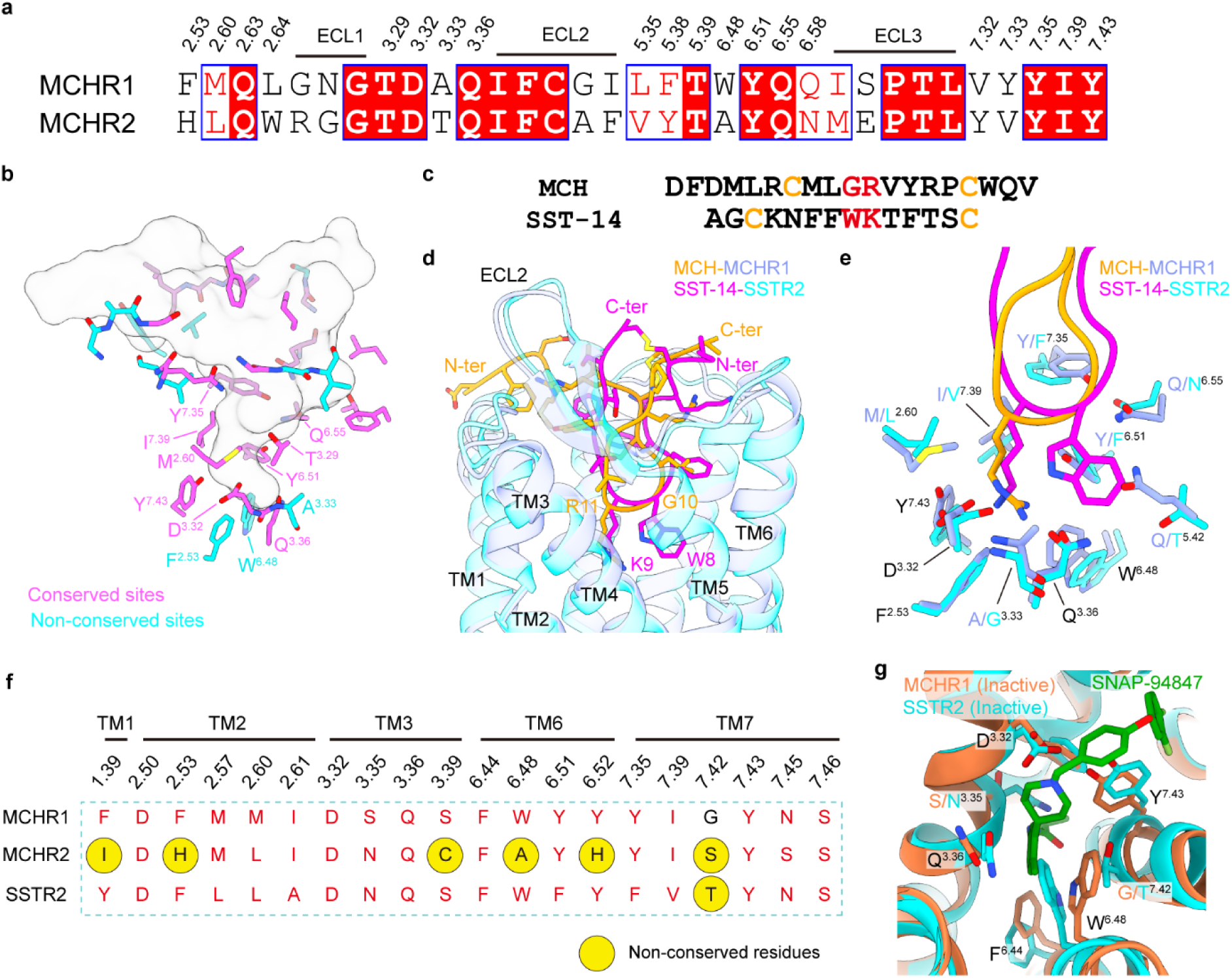
Ligand selectivity of MCHR1. **a**, Sequence alignment of MCH-binding sites in MCHR1 and MCHR2. **b**, Conserved and non-conserved MCH-binding sites in MCHR1 and MCHR2. MCH in the MCH-MCHR1-G_i1_ structure (T1 state) is shown as transparent surface of its atomic model. Residues at conserved sites and non-conserved sites are colored magenta and cyan, respectively. **c**, Sequence alignment of human MCH and SST-14. **d**, Superposition of MCH-MCHR1 (T1 state) and SST-14-bound SSTR1 (PDB ID: 7XMR). **e**, Comparison of the bottom sub-pocket in MCHR1 and SSTR2. **f**, Sequence alignment of SNAP-94847-binding sites in MCHR1, MCHR2, and SSTR2. **g**, Superposition of SNAP-94847-bound MCHR1 and antagonist-bound SSTR2 (PDB ID: 7XNA).

MCHR1 was first found as a somatostatin receptor-like receptor^41^. MCH and somatostatin are both cyclic peptides (Fig. 5c). The binding pocket of MCH in MCHR1 also resembles that of SST-14 in SSTR2^42^. At the bottom of the binding pocket, **R11** of MCH overlaps well with K9 of SST-14, while W8 in SST-14 is replaced by a glycine (**G10**) in MCH (Fig. 5d). The surrounding residues are mostly similar except for a glutamine of MCHR1 at position 5.42, which is substituted by a threonine in SSTR2 for accommodation of the bulky sidechain of W8 (Fig. 5e). However, the top portion of MCH exhibits distinct conformation from SST-14 and binds in a less conserved region (Fig. 5d), which further contributes to ligand specificity. These findings are consistent with the notion that MCHR1 is a specific receptor for MCH that cannot be activated by SST-14^43^.

Many antagonists of MCHR1 have a common 4-arylpiperidine scaffold, which is also present in SNAP-94847. In this study, we revealed the specific binding mode of this molecular feature in MCHR1. The quaternary amine in MCHR1 antagonists is anchored by D192^3.32^. The methylphenyl group connected at the C4 position of the piperidine group tightly packs with W^6.48^ of MCHR1, effectively blocking MCHR1 activation. Docking analysis using the inactive structure suggests that a range of antagonists may share a common mechanism of action (Extended Data Fig. 7).

Our study offers a structural basis for the understanding of previous structure-activity relationship (SAR) studies^34,44^. It was found that unproper substitution on the aryl group of the 4-arylpiperidine scaffold may lead to weakened affinity^34^. From our structure, an apparent explanation is that the current 6-methyl group form optimal hydrophobic interactions with Y341^6.51^ and Y342^6.52^, while a 4-methyl group may sterically clash with F334^6.44^. However, 4,6-diF substitution on this group seems also acceptable for affinity as exemplified by SNAP-102739^44^. In addition, the isopropyl group at the anilide position was reported to dramatically increase the affinity for MCHR1^34^. From the structure, this is perhaps because an isopropyl group is preferred by the small sub-pocket inside the 7TM domain of MCHR1. In the R3 group part, 4-aryloxybenzyl analogues are better than 3-aryloxybenzyl^34^, which can be explained by an optimal steric match with the pocket that accommodates the R3 group. Moreover, small electron-withdrawing groups at the end of the N-alkyl part showed favorable MCHR1 affinities^34^ possibly through interaction with F116^1.39^. These collective effects make SNAP-94847 a high-affinity antagonist of MCHR1. This understanding provides unprecedented opportunities for rational design of better anti-MCHR1 drugs.

Despite the apparent homology between MCHR1 and MCHR2, antagonists inhibit these two receptors differentially. SNAP-7941, an antagonist of MCHR1 with the 4-arylpiperidine scaffold, demonstrates remarkable selectivity for MCHR1 over MCHR2^22^. Through sequence alignment of SNAP-94847-binding sites, we identified several non-conserved sites between the two receptors. Notably, the residue at position 6.48, a tryptophan in MCHR1 that is crucial for interaction with the 4-arylpiperidine scaffold, is replaced by an alanine in MCHR2 (Fig. 5f). This substitution, along with other variations, likely contributes to the selectivity of SNAP-7941 for MCHR1.

In contrast, when aligning the SNAP-94847-interacting residues in MCHR1 with their counterparts in SSTR2, we found that the residues are largely conserved (Fig. 5f), raising the question of why there are no SNAP-94847-like antagonists for SSTR2. The inactive structure of SSTR2 in complex with a peptide antagonist was also reported recently^42^, enabling the analysis of antagonist selectivity. From structural comparison, we reasoned that the binding of the 4-arylpiperidine group of SNAP-94847 in SSTR2 is potentially hindered by T301 at position 7.42, where G369^7.42^ in MCHR1 facilitates the necessary conformational change of W^6.48^ for antagonist binding (Fig. 5g). Besides, although resembling S^3.35^ in MCHR1, N^3.35^ in SSTR2 could potentially clash with the R2 group of SNAP-94847, further influencing ligand selectivity (Fig. 5g).

In summary, our study reveals the molecular basis for hormone and small-molecule antagonist recognition, activation, and G protein coupling of the melanin-concentrating hormone receptor MCHR1. These findings provide insights into MCHR1 signaling and antagonism, as well as determinants of ligand selectivity, laying the groundwork for the development of next-generation MCHR1-targeted drugs.

## Materials and Methods

### Construct generation

For structure determination of the MCH-MCHR1-G_i1_ complex, the wild-type human melanin-concentrating hormone receptor 1 (MCHR1) with a truncated C-terminus (the last 26 amino acids were truncated) was synthesized and constructed into a modified pFastBac1 vector containing a bovine prolactin signal peptide followed by a FLAG tag and an 8× His tag at the N-terminus for purification. To facilitate protein expression, MCHR1 was fused with an N-terminal fragment of β_2_AR (BN). A NanoBiT strategy was applied as previously described^35^. An LgBiT subunit was fused with a 17-amino-acid linker (HMGSSGGGGSGGGGSSG) at the C-terminus of the receptor. Human Gα_i1_ with four dominant-negative mutations (DNGα_i1_)^45^, S47N, G203A, E245A, and A326S, was cloned into the pFastBac1 vector. Human Gβ_1_ with an N-terminal 6× His tag and a C-terminal HiBiT subunit connected by a 15-amino-acid linker (GSSGGGGSGGGGSSG), and human Gγ_2_ were cloned into the pFastBac-Dual vector. Coding sequence of the antibody fragment scFv16 with a GP67 signal peptide at the N-terminus and a TEV cleavage site followed by an 8× His tag at the C-terminus was constructed into the pFastBac1 vector.

For structure determination of the antagonist-bond MCHR1, a previously described strategy was applied^38^. To fuse mBRIL to MCHR1 in a rigid fashion, the active-state structure of MCHR1 was aligned with the previous β_2_AR-mBRIL construct. Appropriate residues were introduced or deleted to obtain a continuous helix for TM5 and TM6. After determining the junction sequences, AlphaFold was used again to predict the resulting sequence to confirm whether a rigid fusion was formed. For the H8 fusion, a similar strategy was used and the coding sequence was constructed into the pFastBac1 vector. For Fab1B3 and the 4-9 glue molecule, the coding sequences were cloned into the pET-22b(+) vector with an N-terminal pelB signal peptide and a C-terminal 6×His tag.

For functional assays, wild-type MCHR1 was cloned into the pcDNA3.1(+) vector before mutations were introduced individually. All constructs were verified by DNA sequencing.

### Protein purification

Expression and purification of scFv16 were conducted as previously described^36^. Briefly, scFv16 was expressed in Tni (HiFive) insect cells and purified by Ni resin. The C-terminal His tag was removed by TEV protease. Proteins were loaded onto a Superdex 200 Increase 100/300 GL column (GE Healthcare) and the correct fractions were pooled, concentrated, flash-frozen and stored at -80 ℃ before use.

For MCH-MCHR1-G_i1_ complex, MCHR1, DNGα_i1_, Gβ_1_, and Gγ_2_ were co-expressed in Sf9 insect cells using the Bac-to-Bac system (Invitrogen). Cells were infected with three types of viruses encoding MCHR1, DNGα_i1_, Gβ_1_γ_2_ at the ratio of 3:2:2 at the density of 2.5 × 10^6^ cells/mL and cultured at 27 ℃ for 48 h. Cells were collected by centrifugation, flash-frozen and stored at -80 ℃ before use. For the purification of MCHR1-G_i1_ complex, cell pellets were thawed in lysis buffer containing 20 mM HEPES pH 7.5, 50 mM NaCl, 10 mM MgCl_2_, 5 mM CaCl_2_, 2.5 μg/mL leupeptin, 300 μg/mL benzamidine, 25 mU/mL Apyrase (New England Biolabs), and 100 μM TCEP at room temperature for 2 h. For MCH-bound complex, MCH peptide (synthesized by Sangon Biotech) was added into the lysis buffer at the final concentration of 2 μM and kept at 1 μM in all the following steps. After centrifugation at 30,700 g for 30 min, the cell membranes were resuspended and solubilized in buffer containing 20 mM HEPES pH 7.5, 100 mM NaCl, 0.5% (w/v) lauryl maltose neopentylglycol (LMNG, Anatrace), 0.1% (w/v) cholesteryl hemisuccinate (CHS, Anatrace), 10% (v/v) glycerol, 10 mM MgCl_2_, 5 mM CaCl_2_, 12.5 mU/mL Apyrase, 2.5 μg/mL leupeptin, 300 μg/mL benzamidine, and 100 μM TCEP for 2 h at 4 ℃. The supernatant was collected by centrifugation at 38,900 g for 45 min and then incubated with Ni resin at 4 ℃ for 2 h. After loaded onto a gravity column, the resin was washed with 20 column volumes of washing buffer containing 20 mM HEPES pH 7.5, 100 mM NaCl, 0.05% (w/v) LMNG, 0.01% (w/v) CHS, 2.5 μg/mL leupeptin, 300 μg/mL benzamidine, 100 μM TCEP, and 20 mM imidazole. Proteins were eluted with the same buffer plus 400 mM imidazole. The eluate was supplemented with 2 mM CaCl_2_ before incubated with anti-FLAG M1 antibody resin overnight at 4 ℃. The FLAG antibody resin was washed with 10 column volumes of washing buffer plus 2 mM CaCl_2_. The complex was eluted with same buffer containing 5 mM EDTA and 200 μg/mL FLAG peptide. Purified scFv16 was added to the eluate at a 1.3:1 molar ratio. Finally, the complex was purified by a Superdex 200 10/300 column (GE Healthcare) equilibrated with buffer containing 20 mM HEPES pH 7.5, 100 mM NaCl, 0.00075% (w/v) LMNG, 0.00025% (w/v) GDN, 0.00015% (w/v) CHS, and 100 μM TCEP. The peak fractions containing monomeric complexes were pooled and concentrated for EM studies.

For Fab1B3 and the glue molecule, the plasmids were transformed into *E.coli* BL21 (DE3) cells, and cells were grown at 37 ℃ in LB medium supplemented with 50 μg/mL ampicillin. Cells were induced by the addition of 1 mM IPTG and incubated for 24 h at 16 ℃. Cells were collected and disrupted in buffer containing 20 mM HEPES pH 7.5 and 150 mM NaCl. Both Fab1B3 and the glue molecule were purified by Ni-affinity chromatography. Unwanted proteins were removed with wash buffer (20 mM HEPES pH 7.5, 150 mM NaCl, and 20 mM imidazole), and the target protein was eluted with wash buffer supplemented with 300 mM imidazole. The eluate was concentrated to 20 mg/ml using a 10 kDa molecular weight cutoff concentrator (Millipore) for the assembly of complexes.

For antagonist-bond MCHR1, the chimeric MCHR1-mBRIL was expressed in Sf9 insect cells using the Bac-to-Bac system. Cells were infected with virus encoding MCHR1-mBRIL at the density of 2.5 × 10^6^ cells/mL and cultured at 27 ℃ for 48 h. Cells were collected by centrifugation, flash-frozen and stored at -80 ℃ before use. Cell pellets were thawed in lysis buffer containing 10 mM HEPES pH 7.5, 0.5mM EDTA, 2.5 μg/mL leupeptin, 150 μg/mL benzamidine at room temperature for 2 h. SNAP-94847 (MedChemExpress) was added into the lysis buffer at the final concentration of 1 μM and kept at 1 μM in all the following steps. Then, excess purified Fab1B3 and the glue molecule were added. After centrifugation at 30,700 g for 30 min, the cell membranes were resuspended and solubilized in buffer containing 20 mM HEPES pH 7.5, 100 mM NaCl, 1% (w/v) LMNG, 0.2% (w/v) CHS, 10% (v/v) glycerol,

2.5 μg/mL leupeptin, 300 μg/mL benzamidine, and 100 μM TCEP for 2 h at 4 ℃. The supernatant was collected by centrifugation at 38,900 g for 60 min and then incubated with Ni resin at 4 ℃ for 2 h. After loaded onto a gravity column, the resin was washed with 20 column volumes of washing buffer containing 20 mM HEPES pH 7.5, 150 mM NaCl, 0.05% (w/v) LMNG, 0.01% (w/v) CHS, 2.5 μg/mL leupeptin, 300 μg/mL benzamidine, 100 μM TCEP, and 20 mM imidazole. Proteins were eluted with the same buffer plus 400 mM imidazole. The eluate was supplemented with 5 mM CaCl_2_ before incubated with anti-FLAG M1 antibody resin overnight at 4 ℃. The resin was washed with 10 column volumes of washing buffer plus 2 mM CaCl_2_. The complex was eluted with same buffer containing 5 mM EDTA and 200 μg/mL FLAG peptide. Finally, the complex was purified by a Superdex 200 10/300 column equilibrated with buffer containing 20 mM HEPES pH 7.5, 150 mM NaCl, 0.00075% (w/v) LMNG, 0.00025% (w/v) GDN, 0.00015% (w/v) CHS, and 100 μM TCEP. The peak fractions containing monomeric complexes were pooled and concentrated for EM studies.

### Cryo-EM sample preparation and data acquisition

An aliquot of 3 μL MCH-MCHR1-G_i1_ complex at the concentration of 5 mg/mL or 3 μL SNAP-94847-bound MCHR1-mBRIL-Fab1B3-Glue complex at the concentration of 3 mg/mL was applied to a glow-discharged holey Ni-Ti alloy grid (ANTcryo, M01, Au300 R1.2/1.3). The grid was blotted and frozen in liquid ethane using Vitrobot Mark IV (Thermo Fischer Scientific). The grids were imaged on a 300 kV Titan Krios electron microscope (Thermo Fischer Scientific) equipped with Gatan K3 Summit direct electron detector and an energy filter. Data were collected at the magnification of 81,000× at a pixel size of 0.535 Å in super-resolution mode using the EPU software. Image stacks were recorded in 32 frames at a total dose of 55 e^-^/Å^2^ with the defocus range from -2.2 to -1.2 μm. A total of 4,901 movies for MCH-MCHR1-G_i1_ complex and 5713 movies for SNAP-94847-bound MCHR1-mBRIL-Fab1B3-Glue complex were collected.

### Cryo-EM data processing

For MCH-MCHR1-G_i1_ complex, 4,901 movies were subjected to CryoSPARC^46^ and processed with Patch motion correction and Patch CTF estimation. Exposures with tolerable CTF fit resolution and total motion distance were selected for further processing. Blob picker was used to pick particles from a small subset of micrographs for creation of 2D templates. Particles were picked from the whole dataset by Template picker using these 2D templates. After 2D classification, six 3D classes were generated by Ab-initio Reconstruction. Particles from 4 classes were further classified by Heterogeneous Refinement. Then two classes with acceptable quality were pooled and re-classified into 6 classes using Ab-initio Reconstruction and Heterogeneous Refinement. Four classes with 871,951, 771,833, 588,396, and 500,329 particles were individually processed by Non-uniform Refinement^47^ and improved by Local Refinement with a customized global mask. The final maps reach the nominal resolution of 2.61 Å, 2.65 Å, 2.78 Å, and 2.81 Å at a Fourier shell correlation (FSC) threshold of 0.143. Estimation of local resolution and local filtering of the maps were performed in CryoSPARC.

For SNAP-94847-MCHR1-mBRIL complex, 5713 movies were subjected to CryoSPARC and processed with MotionCor2 and CTFFIND4. Exposures with tolerable CTF fit resolution and total motion distance were selected for further processing. Blob picker was used to pick particles from a small subset of micrographs for creation of 2D templates. Particles were picked from the whole dataset by Template picker using these 2D templates. After 2D classification, four 3D classes were generated by Ab-initio Reconstruction. Then all particles were classified into these four classes by Heterogeneous Refinement. Three classes with acceptable quality were pooled and re-classified into 4 classes using Ab-initio Reconstruction and Heterogeneous Refinement. Two classes with 305,549 and 268,193 particles were individually processed by Non-uniform Refinement. The final maps reached the nominal resolution of 3.33 Å and 3.43 Å at a Fourier shell correlation (FSC) threshold of 0.143. Estimation of local resolution of the maps were performed in CryoSPARC.

### Model building and validation

For MCH-MCHR1-G_i1_-scFv16 complex, the initial model of MCHR1 was generated by Alphafold^48^. Coordinates of G_i1_-scFv16 was derived from the μOR-G_i1_ complex (PDB ID: 6DDE). MCH was manually built in Coot according to the density.

For SNAP-94847-MCHR1-mBRIL-Fab1B3-Glue complex, the initial model of MCHR1-mBRIL was also generated by Alphafold. Coordinates of mBRIL and Fab1B3 was derived from the crystal structure of BRIL in complex with Fab1B3 (PDB ID: 8J7E). Coordinates of E3 and K3 helices were derived from the structure of the E3/K3 coiled-coil (PDB ID: 1U0I). Coordinates of NbFab were derived from an NbFab-contained cryo-EM structure (PDB ID: 7PHP). Coordinates of ALFA tag and NbALFA were derived from the crystal structure of NbALFA bound to ALFA tag peptide (PDB ID: 6I2G). Coordinates and geometry restrains of SNAP-94847 were generated using eLBOW in Phenix.

The models were fitted into the EM map and combined using UCSF Chimera^49^. Then the model was corrected by manual adjustment in Coot^50^ and refined by Real-space refinement in Phenix^51^. Model statistics were calculated by MolProbity^52^ and provided in Supplementary Table 1.

### G protein-dissociation assay

Function of wild-type and mutant MCHR1 was measured using the TRUPATH biosensors as previously described^53^. HEK293T cells were distributed into six-well plates at a density of 1.2 × 10^6^ cells per well and incubated for 8 h at 37 ℃. A plasmid mixture of 0.5 μg wild-type or mutant MCHR1, 0.5 μg Gα_i1_-RLuc8, 0.5 μg Gβ_3_, 0.5 μg GFP2-Gγ_9_ was co-transfected into HEK293T cells using Lipofectamine 3000 (Thermo Fisher Scientific). After 40 h, cells were harvested, washed with HBSS (Hank’s Balanced Salt Solution), and resuspended in 800 μL BRET buffer (HBSS supplemented with 25 mM HEPES pH 7.4 and 0.1% BSA). Cells were divided into white-wall white-bottom 96-well plates at the density of 100,000 cells per well. Then the luciferase substrate coelenterazine 400a at 5 μM working concentration was added and the plates were incubated at room temperature for 5 minutes. Cells were stimulated with MCH at different final concentrations before the plates were incubated for another 5 minutes at room temperature. The BRET signal was recorded by SpectraMax iD5 (Molecular Devices) and calculated as the ratio of light emission at 515 nm (GFP2)/410 nm (RLuc8). Data were baseline-corrected with the ligand-free control and curves were calculated by a three-parameter logistic function. Data from three independent experiments were used for analysis.

To measure the activity of antagonist SNAP-94847, we carried out the same procedures as those for MCHR1-mediated Gα_i1_ dissociation from Gβ_1_γ_2_, except that HEK293T cells were pre-treated with different concentrations of SNAP-94847 dissolved in assay buffer from 10^−1^^1^ M to 10^−4^ M and incubated for 10 min. After that, 10 μM MCH were added to the wells and incubated for 5 min. The BRET signal was recorded by SpectraMax iD5 (Molecular Devices) and calculated as the ratio of light emission at 515 nm (GFP2)/410 nm (RLuc8). Data were baseline-corrected with the antagonist-free control and curves were calculated by a three-parameter logistic function. Data from three independent experiments were used for analysis.

### Cell-surface expression analysis

Cell-surface expression of wild-type MCHR1 and mutants was measured by a fluorescence-activated cell sorting (FACS) assay. HEK293T cells were seeded in 24-well plates at the density of 2 × 10^5^ cells per well before transfected with 0.5 μg plasmid encoding FLAG-tagged wild-type MCHR1 or mutants using Lipofectamine 3000. After 42 h, cells were collected and resuspended in HBSS. 20 μL cells were incubated with 20 μL anti-FLAG M2-FITC antibody (Sigma Aldrich) diluted in TBS buffer containing 20 mM Tris pH 7.5, 150 mM NaCl, and 4% (w/v) BSA at 4 ℃ for 20 min. 160 μL HBSS buffer supplemented with 5 mM HEPES pH 7.4 was added after incubation. The fluorescence was measured on CytoFLEX (Beckman). The gate was set by FSC/SSC thresholds to define single cells. Surface expression level was evaluated by mean fluorescence intensity and normalized to the mock and wild-type group. Data from three independent experiments were used for analysis.

### Molecular docking

To investigate the binding modes of other antagonists of MCHR1, we performed molecular docking studies using AutoDock Vina^54^. MCHR1 from the cryo-EM structure of SNAP-94847-MCHR1 was used as the receptor. Coordinates of different antagonists were downloaded from PubChem. Coordinates of the receptor and the ligands were processed by AutoDockTools using default settings. The docking box was a 15∼18 Å cube centered on SNAP-94847 in the cryo-EM structure. No flexible residues of the receptor were defined. Binding poses were selected according to binding energy and visual inspection.

## Acknowledgements

We thank the Cryo-EM Center at University of Science and Technology of China for the support of cryo-EM data collection. We thank Dr. Yongxiang Gao for assistance with cryo-EM data screening and collection. This work is supported by the Ministry of Science and Technology of China (grant number 2019YFA0904100 and 2021YFA0910202), the Strategic Priority Research Program of the Chinese Academy of Science (Grant No. XDB0540000) and Natural Science Foundation of China (grant number T2221005).

## Author contributions

W.G. and H.L. conceived the study. X.Y. expressed and purified the protein complexes with constructs from X.L., B.H., and Y.T. X.Y. and G.L. prepared the grids, collected and processed cryo-EM data, built the models, and analyzed the structures. X.Y. performed the functional assays and analyzed the results. X.Y. and G.L. wrote the manuscript under the supervision of H.L. and W.G.

## Data availability

The atomic coordinates of MCH-MCHR1-G_i1_ complex in different states (T1, T2, L1, and L2) have been deposited in the Protein Data Bank (PDB) under accession codes xxxx, xxxx, xxxx, and xxxx, respectively. The EM maps of MCH-MCHR1-G_i1_ complex have been deposited in the Electron Microscopy Data Bank (EMBD) under accession codes EMD-xxxxx, EMD-xxxxx, EMD-xxxxx, and EMD-xxxxx, respectively. The atomic coordinates of SNAP-79847-bound MCHR1-mBRIL complex in S1 and S2 states have been deposited in the Protein Data Bank (PDB) under accession codes xxxx and xxxx, respectively. The EM maps of SNAP-79847-bound MCHR1-mBRIL complex have been deposited in the Electron Microscopy Data Bank (EMBD) under accession codes EMD-xxxxx and EMD-xxxxx, respectively.

