## Supplementary materials for "Structural insights into physiological activation and antagonism of melanin-concentrating hormone receptor MCHR1"

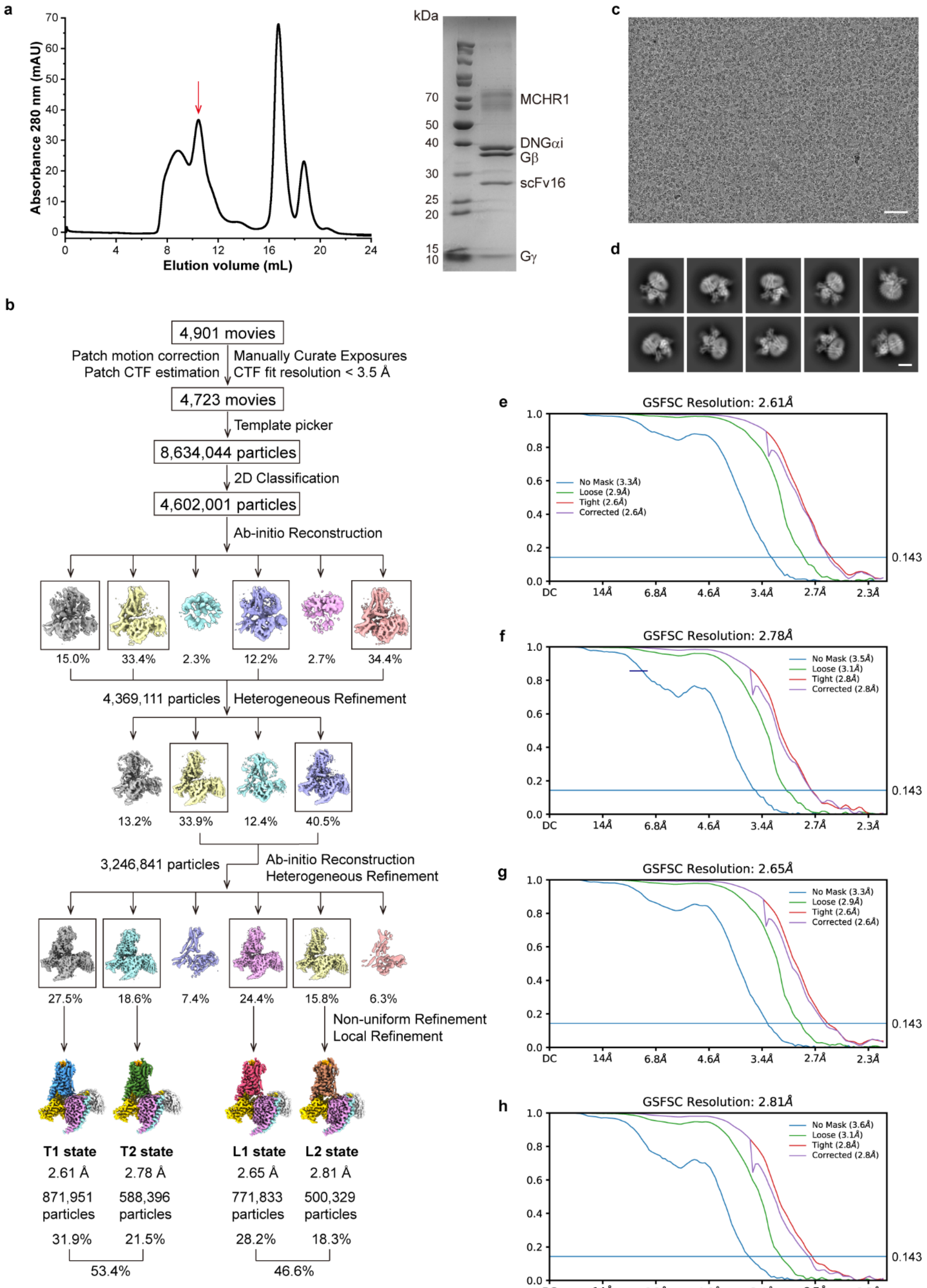


**Extended Data Fig. 1 | Structure determination of MCH-MCHR1-$G_{i1}$ complex by cryo-EM. a,**

Size-exclusion chromatography (SEC) and SDS-PAGE (stained by Coomassie blue) profiles of the MCH-bound MCHR1-$G_{i1}$ complex. **b**, Processing workflow of cryo-EM data. **c**, Representative micrograph. **d**, Representative 2D averages. **e-h**, Gold-standard FSC curves for EM maps of T1 state, T2 state, L1 state, and L2 state, respectively.

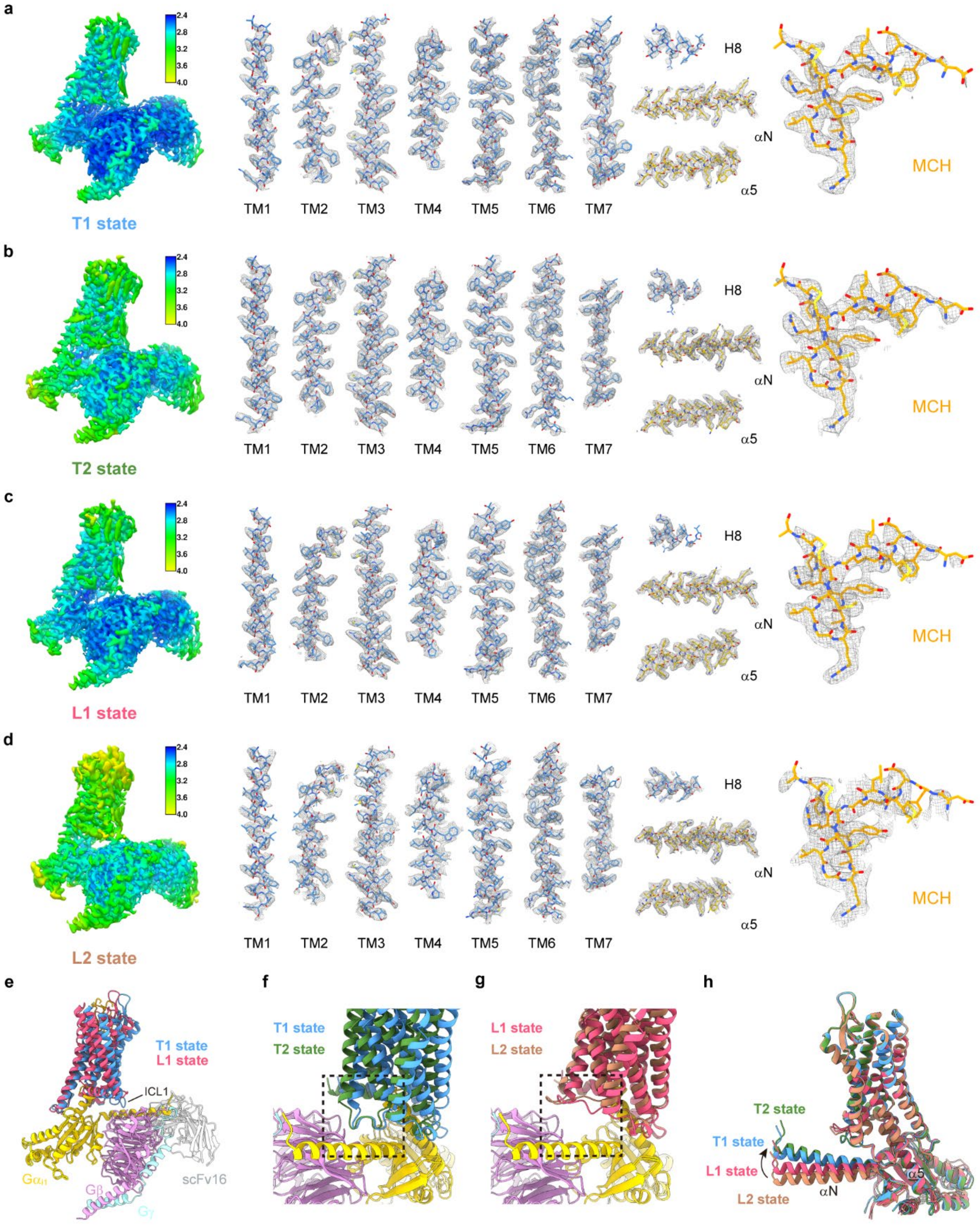


**Extended Data Fig. 2 | Local resolution, density maps and differences of MCH-MCHR1-$G_{i1}$ structures. a-d**, Local resolution of EM maps and density maps of transmembrane helices (TM1-TM7), H8, αN helix, α5 helix and MCH. EM maps and density maps are displayed at the contour level of 0.40, 0.33, 0.35, and 0.23 for T1 state, T2 state, L1 state, and L2 state, respectively. **e**, Superposition of T1 state and L1 state (aligned by Gβ). **f-g**, The contact between ICL1 and Gi1 in the tight conformers (**f**) or the loose conformers (**g**). **h**, The rotation of αN helix in 4 states (aligned by MCHR1).

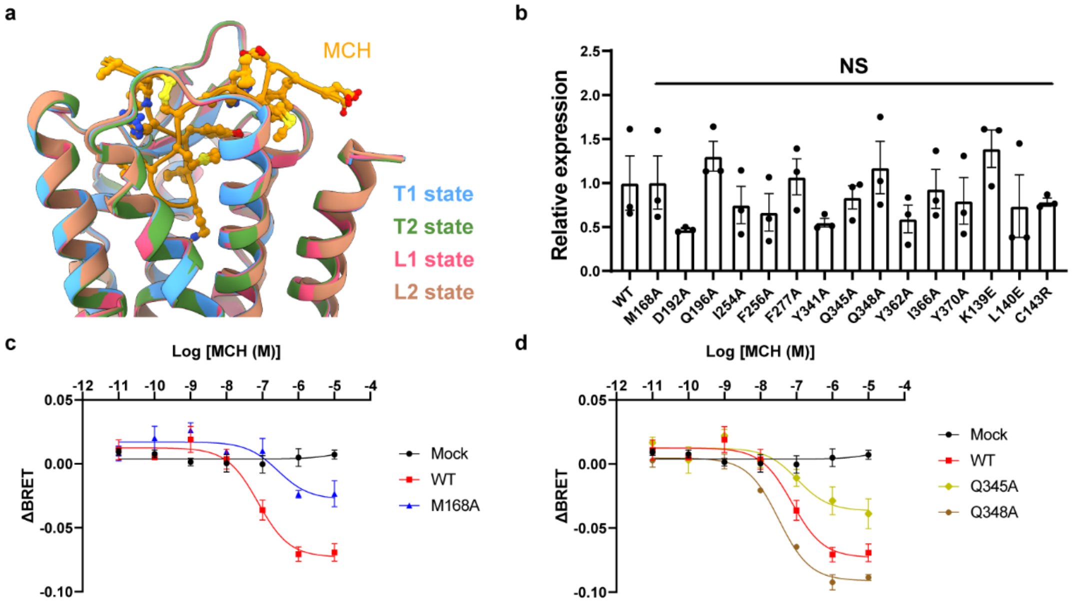


**Extended Data Fig. 3 | Structural comparison and functional data for the active structure. a**, Superposition of different conformers of the MCH-MCHR1-$G_{i1}$ complex. **b**, Relative cell-surface expression of MCHR1 mutants in HEK293T cells. Data are shown as means ± SEM from three independent experiments. Expression of each mutant is compared to WT using one-way ANOVA with Dunnett's multiple comparisons. NS, no significant difference. **c**, $G_i$-dissociation curve of M168A mutant. **d**, $G_i$-dissociation curves of Q345A and Q348A mutants.

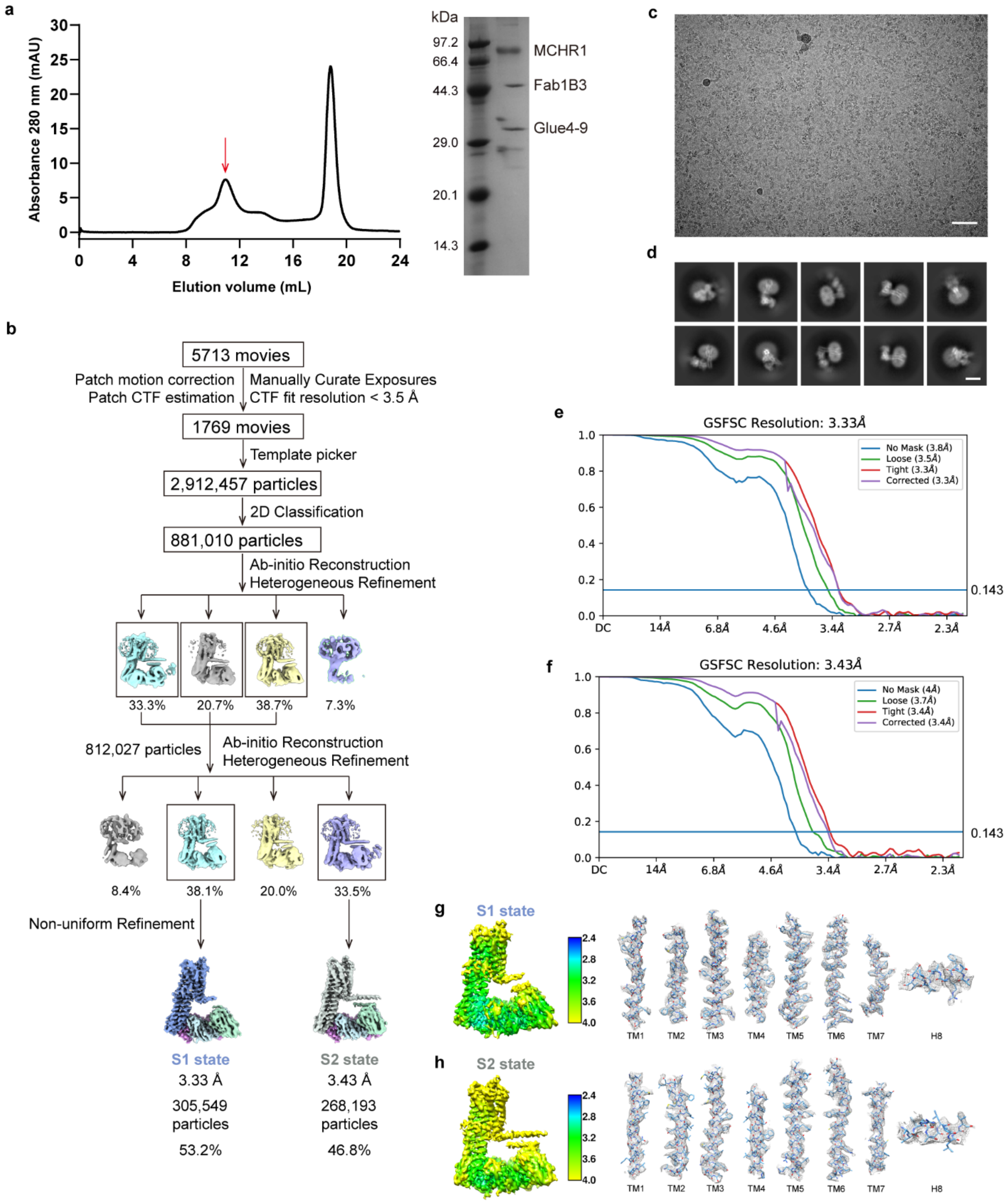


**Extended Data Fig. 4 | Structure determination of antagonist-bound MCHR1 by cryo-EM. a**, Size-exclusion chromatography (SEC) and SDS-PAGE (stained by Coomassie blue) profiles of the MCHR1-Fab1B3-Glue complex. **b**, Processing workflow of cryo-EM data. **c**, Representative micrograph. **d**, Representative 2D averages. **e-f**, Gold-standard FSC curves for EM maps of S1 state and S2 state, respectively. **g-h**, Local resolution of EM maps and density maps of transmembrane helices (TM1-TM7) and H8. EM maps and density maps are displayed at the contour level of 0.45 and 0.40 for S1 state and S2 state, respectively.

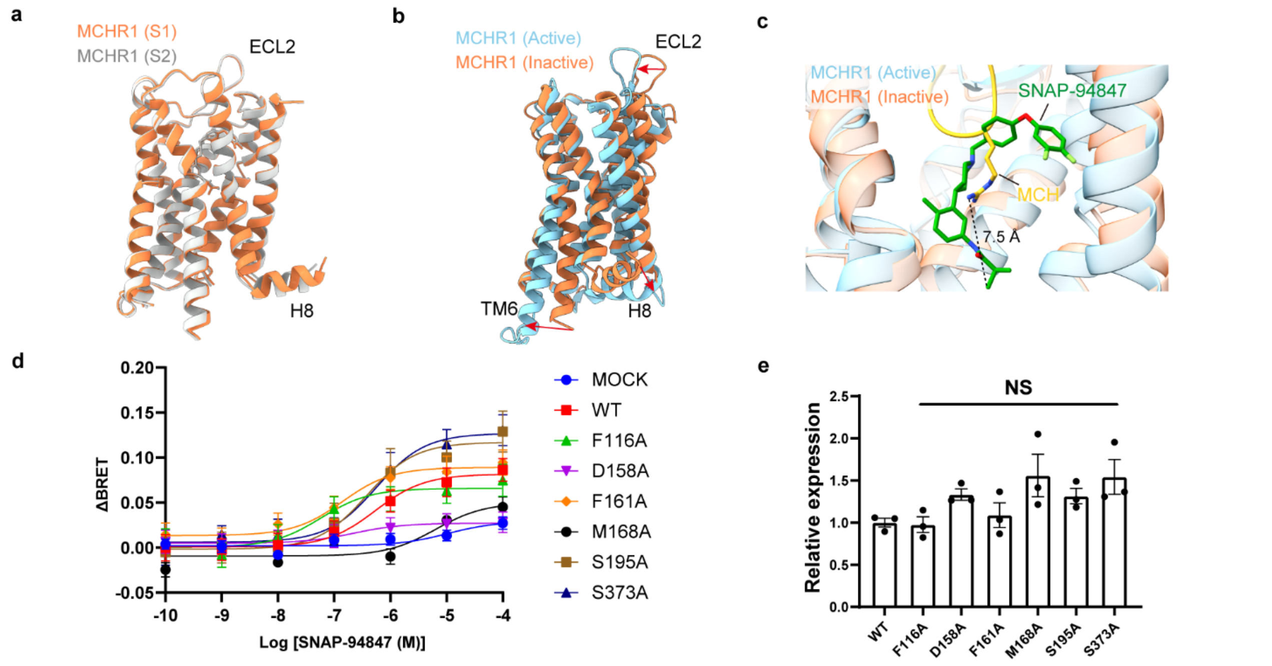


**Extended Data Fig. 5 | Structural comparison and functional data for the inactive structure.** **a**, Comparison of S1 state and S2 state of antagonist-bound MCHR1. **b**, Superposition of the active (T1 state) and the inactive (S1 state) structures of MCHR1. **c**, Comparison of the binding pockets for MCH and SNAP-79847. **d**, $G_i$-dissociation curves of MCHR1 mutants in response to SNAP-94847. Data are shown as means ± SEM from three independent experiments. **e**, Relative cell-surface expression of MCHR1 mutants in HEK293T cells. Data are shown as means ± SEM from three independent experiments. Expression of each mutant is compared to WT using one-way ANOVA with Dunnett's multiple comparisons. NS, no significant difference.

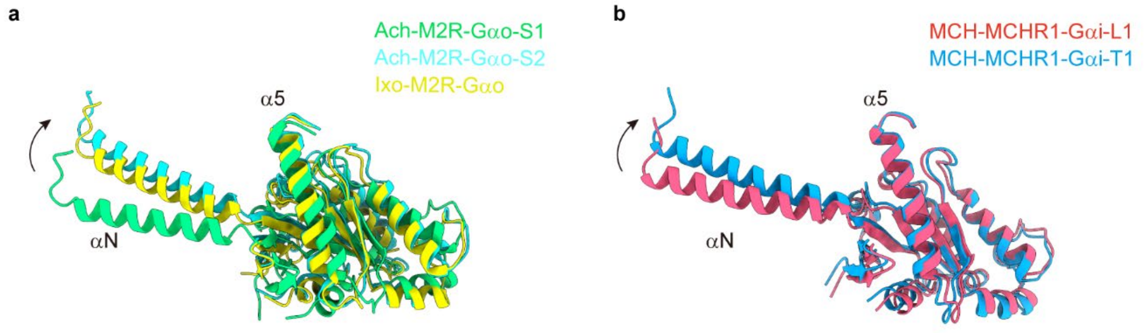


**Extended Data Fig. 6 | Comparison of the orientation of Gα. a**, Comparison of the orientation of $G\alpha_o$ in ACh-bound S1 (PDB ID: 7T8X), ACh-bound S2 (PDB ID: 7T90), and Ixo-bound (PDB ID: 6OIK) states. **b**, Comparison of the orientation of $G\alpha_i$ in MCH-bound L1 and T1 states.

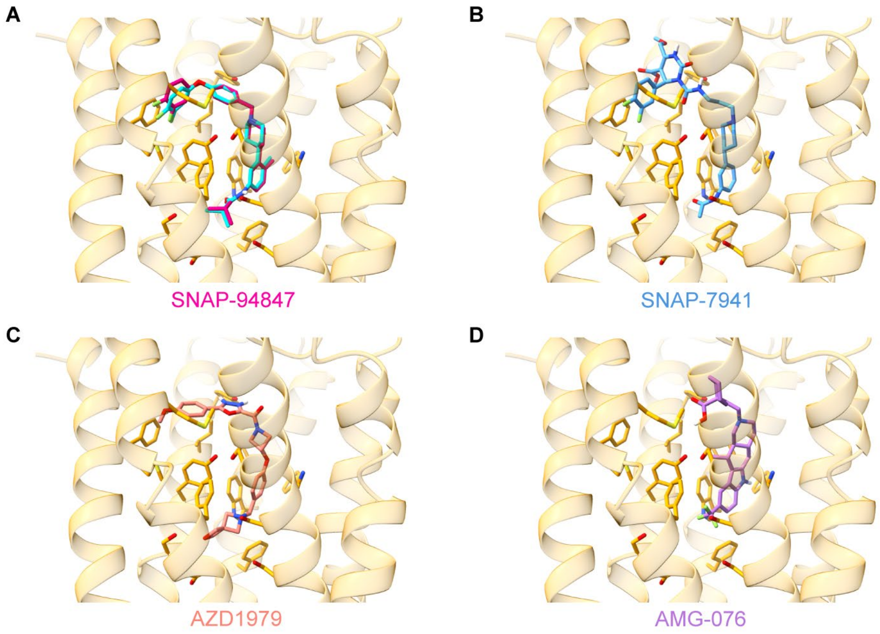


**Extended Data Fig. 7 | Docking results of MCHR1 antagonists. a**, Superposition of SNAP-94847 in the cryo-EM structure (cyan) and the docked pose (magenta). **b-d**, Docked poses of other MCHR1 antagonists.

### Supplementary Table 1 | Cryo-EM data collection, refinement and validation statistics

| | **MCH-MCHR1-$G_{i1}$ T1 state (EMD-77777) (PDB XXXX)** | **MCH-MCHR1-$G_{i1}$ T2 state (EMD-77777) (PDB XXXX)** | **MCH-MCHR1-$G_{i1}$ L1 state (EMD-77777) (PDB XXXX)** | **MCH-MCHR1-$G_{i1}$ L2 state (EMD-77777) (PDB XXXX)** | **SNAP-94847-MCHR1 S1 state (EMD-77777) (PDB XXXX)** | **SNAP-94847-MCHR1 S2 state (EMD-77777) (PDB XXXX)** |
|---|---|---|---|---|---|---|
| Magnification | 81,000 | 81,000 | 81,000 | 81,000 | 81,000 | 81,000 |
| Voltage (kV) | 300 | 300 | 300 | 300 | 300 | 300 |
| Electron exposure ($e^-/Å^2$) | 55 | 55 | 55 | 55 | 55 | 55 |
| Defocus range (μm) | -2.2 to -1.2 | -2.2 to -1.2 | -2.2 to -1.2 | -2.2 to -1.2 | -2.2 to -1.2 | -2.2 to -1.2 |
| Pixel size (Å) | 1.07 | 1.07 | 1.07 | 1.07 | 1.07 | 1.07 |
| Symmetry imposed | C1 | C1 | C1 | C1 | C1 | C1 |
| Initial particle images (no.) | 8,634,044 | 8,634,044 | 8,634,044 | 8,634,044 | 2,912,457 | 2,912,457 |
| Final particle images (no.) | 871,951 | 588,396 | 771,833 | 500,329 | 305,549 | 268,193 |
| Map resolution (Å) | 2.61 | 2.78 | 2.65 | 2.81 | 3.33 | 3.43 |
| FSC threshold | 0.143 | 0.143 | 0.143 | 0.143 | 0.143 | 0.143 |
| Map resolution range (Å) | 2.5-5.0 | 2.5-5.0 | 2.5-5.0 | 2.5-5.0 | 2.5-5.0 | 2.5-5.0 |
| **Refinement** | | | | | | |
| Initial model used (PDB code) | 6DDE | 6DDE | 6DDE | 6DDE | - | - |
| Model resolution (Å) | 2.71 | 2.87 | 2.75 | 2.96 | 3.42 | 3.59 |
| FSC threshold | 0.5 | 0.5 | 0.5 | 0.5 | 0.5 | 0.5 |
| Model composition | | | | | | |
| Non-hydrogen atoms | 8967 | 8964 | 8960 | 8963 | 7296 | 7420 |
| Protein residues | 1160 | 1160 | 1160 | 1160 | 958 | 974 |
| Ligands | 0 | 0 | 0 | 0 | 1 | 1 |
| B factors ($Å^2$) | | | | | | |
| Protein | 32.30 | 39.99 | 30.12 | 62.41 | 89.47 | 86.85 |
| Ligand | - | - | - | - | 96.64 | 106.19 |
| R.m.s. deviations | | | | | | |
| Bond lengths (Å) | 0.002 | 0.002 | 0.002 | 0.004 | 0.002 | 0.002 |
| Bond angles (°) | 0.392 | 0.438 | 0.464 | 0.501 | 0.396 | 0.389 |
| Validation | | | | | | |
| MolProbity score | 1.25 | 1.32 | 1.26 | 1.32 | 1.23 | 1.58 |
| Clashscore | 4.78 | 5.86 | 4.96 | 5.91 | 3.96 | 5.25 |
| Poor Rotamers (%) | 0.72 | 0.83 | 0.62 | 0.93 | 1.16 | 1.65 |
| Ramachandran plot | | | | | | |
| Favored (%) | 99.21 | 98.43 | 98.78 | 98.25 | 98.42 | 97.30 |
| Allowed (%) | 0.79 | 1.57 | 1.22 | 1.75 | 1.58 | 2.70 |
| Disallowed (%) | 0 | 0 | 0 | 0 | 0 | 0 |
